# A Benchmarking Study of Feature Screening Approaches Across Omics Classification Settings

**DOI:** 10.64898/2026.02.25.706632

**Authors:** Erik D. VonKaenel, Lisa M. Bramer, Javier E. Flores, Thomas O Metz, Ernesto S. Nakayasu, Bobbie-Jo M. Webb-Robertson

## Abstract

In recent years, high dimensional omics analyses have become more commonplace for investigating complex biological systems. Typically, these studies attempt to identify key biomolecules associated with a particular biological process. Often, machine learning (ML) is used to identify these biomolecules, typically by learning which biomolecules are highly predictive of a treatment, biological outcome, or phenotype. A major challenge of applying ML to high throughput omics is overcoming noise when sample size is limited and unbalanced with respect to tens of thousands of biomolecules measured. Thus, feature selection (the process of reducing the number of predictors) is both a critical and common step in the ML analysis pipeline. While much attention has been given to embedding and wrapping techniques for feature selection in the omics space, filter-based methods for model-free feature selection have appealing theoretical properties. This manuscript evaluates sure screening, a class of filter-based feature selection methods which provide analytical guarantees for true feature set retention. Here, we cover existing feature screening methods based on the sure screening principal, available software, methods to improve feature screening, and contextualize feature screening in the larger discussion of feature selection for omics data analysis. Additionally, a suite of model-free sure screening approaches is applied and compared for several omics biomedical applications in a ML classification context. We identified BcorSIS as the most effective and computationally efficient screening method across various omics datasets, consistently outperforming others like CSIS and DCSIS in runtime.

**Author Summary:** A common goal when analyzing high throughput omics data is to identify key biomolecules which may be important in complex biological systems. This type of large-scale analysis is critical, as it can significantly reduce the set of total biomolecules which are considered for more detailed, targeted analysis. Additionally, biomolecules which are highly predictive of specific phenotypes are useful markers to guide treatment. For instance, identifying which biomolecules are predictive of type 1 diabetes could improve early diagnosis, which may improve patient prognosis. However, modern instrumentation can often detect thousands or tens of thousands of biomolecules, and studies typically have a disproportionately limited number of samples. Many measured biomolecules, or features, are often noisy and uninformative, and overcoming this noise to identify an informative set of biomolecules can be challenging for machine learning (ML) models. Fortunately, there are many feature selection strategies which can intelligently reduce the number of “uninformative” biomolecules an ML model needs to overcome. In this work, sure screening, which is a class of feature selection methods that retains the true feature set under certain conditions, is evaluated in the context of omics data analyses. The pros and cons of various sure screening methods, including software availability, are summarized, and their use in the larger feature selection literature is contextualized. Additionally, we benchmark performance by applying a suite of sure screening approaches on several real omics datasets generated to understand the progression of type 1 diabetes.

## Introduction

High throughput omics technologies are popular for the investigation of biological systems. Similarly, it’s become commonplace to use machine learning (ML) as a tool for identifying key biomarkers within these datasets and which are highly predictive of a disease^1–4^. However, the number of biomolecules measured by modern omics technologies is as high as tens-of-thousands (e.g. transcriptomics^5^, post-translational modifications^6,7^). Despite the large number of observations, many of these biomolecules are irrelevant to the specific biological question associated with the study. Thus, feature selection is often performed in conjunction with predictive models and analyses such as ML to remove these non-informative features.

In existing literature, feature selection methods have been separated into three primary categories, namely, filters, wrappers, and embedders^8–12^. Filters can generally be thought of as any feature selection method, independent of the model, that involves determining some threshold to retain or remove predictors based on a defined metric. Filter methods can be further categorized as univariate or multivariate. Univariate methods rely on some pairwise similarity metric between each feature and the response, while multivariate methods incorporate feature dependencies. Wrappers can be thought of as methods “wrapping” the feature selection process around a classifier. An example of this is simulated annealing^13^ or repeated optimization for feature interpretation (ROFI)^14^, where a classifier is trained at each step and a feature selection procedure systematically adds or removes features from the model based on the performance of the classifier. An embedder is any method which includes feature selection as a part of the model’s optimization process. Common examples of embedders are random forests^15^ and penalized generalized linear models^16^.

There is often a trade-off when determining an appropriate feature selection strategy. Commonly used filter based methods are generally easier to implement with low computational complexity but can potentially result in feature sets with lower predictive performance than other approaches^12,17^. Generally, wrappers fall on the other end of the spectrum, potentially yielding feature sets highly predictive of the response but with a much larger computational cost, while embedders typically fall somewhere in between filter and wrapper methods with respect to both computational cost and performance^12,17^. This trade-off may seem to be an all-or-nothing choice between methods. However, it may often be appropriate to consider a multi-stage feature selection procedure, such as first applying a filter method, followed by an embedder or wrapper. Many developers in the feature screening space advocate for multi-stage methods to capitalize on the benefits of many approaches^18–20^. Yet, common filter methods are often overlooked, particularly in reviews, due to their simplicity and historically strict assumptions about the data generating mechanism, which may not be applicable to omics data types.

While several reviews of feature selection methods exist, both in general^8,12,17,21,22^ and in omics applications^9–11,23^, there is often a heavy emphasis on wrappers and embedders, while limiting the scope of filters to simple approaches and heuristics which may not be the most appropriate for omics classification contexts. For instance, Lualdi et al^10^ review feature selection methods in proteomics applications, but only discuss simple similarity metrics such as correlation and information gain, in addition to routine approaches for handling multiple comparisons. Li et al^11^ benchmark four filter methods, two wrappers, and two embedding approaches across fifteen multi-omics cancer datasets, though the filter methods are limited to a basic t-test, information gain, and two classical algorithms: minimum redundance maximum relevancy (mRMR) and reliefF. Similarly, Abusamra et al^9^ evaluated feature selection in gene expression data, but the filter methods are also limited to classical approaches such as t-tests and information gain. While these approaches are computationally efficient, they often make assumptions that are too strict about the data generating mechanism, do not provide analytic guarantees about feature retention, or rely on a heuristically determined cutoff.

In this manuscript, we focus on more sophisticated filter methods with minimal assumptions about the data generating mechanism and analytic guarantees regarding feature retention. A particularly rich base of literature in this space is sure screening, which remain relatively isolated from feature selection in omics ML applications. Sure screening has been reviewed occasionally in the past, with the first instance we are aware of presented in Fan et al^24^ mentioning independence screening as a small part of a larger review on the challenges of big data. The authors of the original sure screening method provide an extensive review of sure independence screening as of 2018^25^, but do not approach the review from an omics application perspective. Furthermore, additional methods have been developed since the release of the paper, and they do not discuss software implementations of the methods. Lastly, the He et al^26^ review covers sure screening methods for multivariate responses, and the Liu et al^27^ review on feature selection mentions sure screening, but neither offers any discussion regarding omics data analysis applications. In this manuscript, rather than focus on the theoretical specifics of any single screening method, a general overview of the topic for omics practitioners is provided, discussing high level pros and cons of different approaches including software availability. Further, a suite of sure screening methods is applied to several real data applications in the omics space to evaluate utility in practice.

## Screening Methods

Without loss of generality, the feature screening problem can be formalized in a classical linear regression setting. In a standard high dimensional linear model

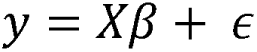

where *y*= (*Y* _1_,…,*Y_n_*)*^T^* represents the outcome, *X*= (*x*_1_,…,x*_n_*)*^T^* are the biomolecules, *ϵ* = (*ϵ*_1_,…,*ϵ_n_*)*^T^* is an additive noise term, {*β* = ({*β_1_*,…, *β_p_*)*^T^* are the covariates, and *p*> *n*. Some amount of sparsity is often assumed in the feature space, i.e., {*β* is sparse. In the context of omics, this equates to only a subset of measured biomolecules having some (linear) relationship with the response. To improve, the predictive performance or runtime of machine learning methods, one can attempt to screen, or remove, features which are unrelated to the response. In this context, when feature screening is discussed, this is referring to some feature selection procedure which only involves computing a model independent measure of association. For example, in the case of standard linear regression, independence screening might be operationalized as performing a series of marginal t-tests and only keeping those features with a p-value below some pre-specified cutoff. Compared to alternative methods of feature selection, such as wrapper and embedding methods, feature screening is often more computationally efficient and easily scalable to ultra-high dimensional datasets where the feature set grows at a super polynomial rate with respect to the sample size. A primary challenge when using a screening strategy is ensuring signal is retained while maintaining some amount of robustness to both the data generative mechanism, and the predictive model being used. For instance, in the context of omics, a screening strategy should allow for potentially complicated non-linear relationships between the features and the response. Over the past decade, several “model-free” screening metrics have been proposed, significantly relaxing assumptions about the model and data. The bulk of this work focuses on model-free screening procedures, as these methods are the most relevant to omics applications. In the following sections, classical approaches along with recent advances in screening methods, which can have the potential to tackle the challenges associated with omics data and ML, are discussed.

### Sure Screening

A now foundational concept in the screening literature is “sure screening”. This term was introduced as a property that any screening procedure can possess^20^. Specifically, a screening procedure has the sure screening property if all “important variables survive … with probability tending to 1”. This concept was formalized by showing that Pearson correlation has the sure screening property under the assumptions of the classic linear model^20^. A notable caveat of sure screening is that the selection of important features is shown to be asymptotic with sample size. Additionally, it has been shown that there exists a relationship between the model size which retains all important features, and both the signal in the data and collinearity among features ^20^. The lower the signal and/or the more collinearity, the larger the selected model needs to be to retain the “important” feature set.

Since the work of Fan et al^20^, many have proposed new measures of association that have the sure screening property, with progressively relaxed assumptions about the underlying model and data generating mechanisms. These methods all generally share similar assumptions: there exists some model size which asymptotically retains the important feature set given enough signal in the data. However, various metrics respond differently depending on the levels of noise and collinearity. The key aspects of each screening method are given in Table 1. As noted in Table 1, most of these methods are available in open-source software.

**Table 1:**
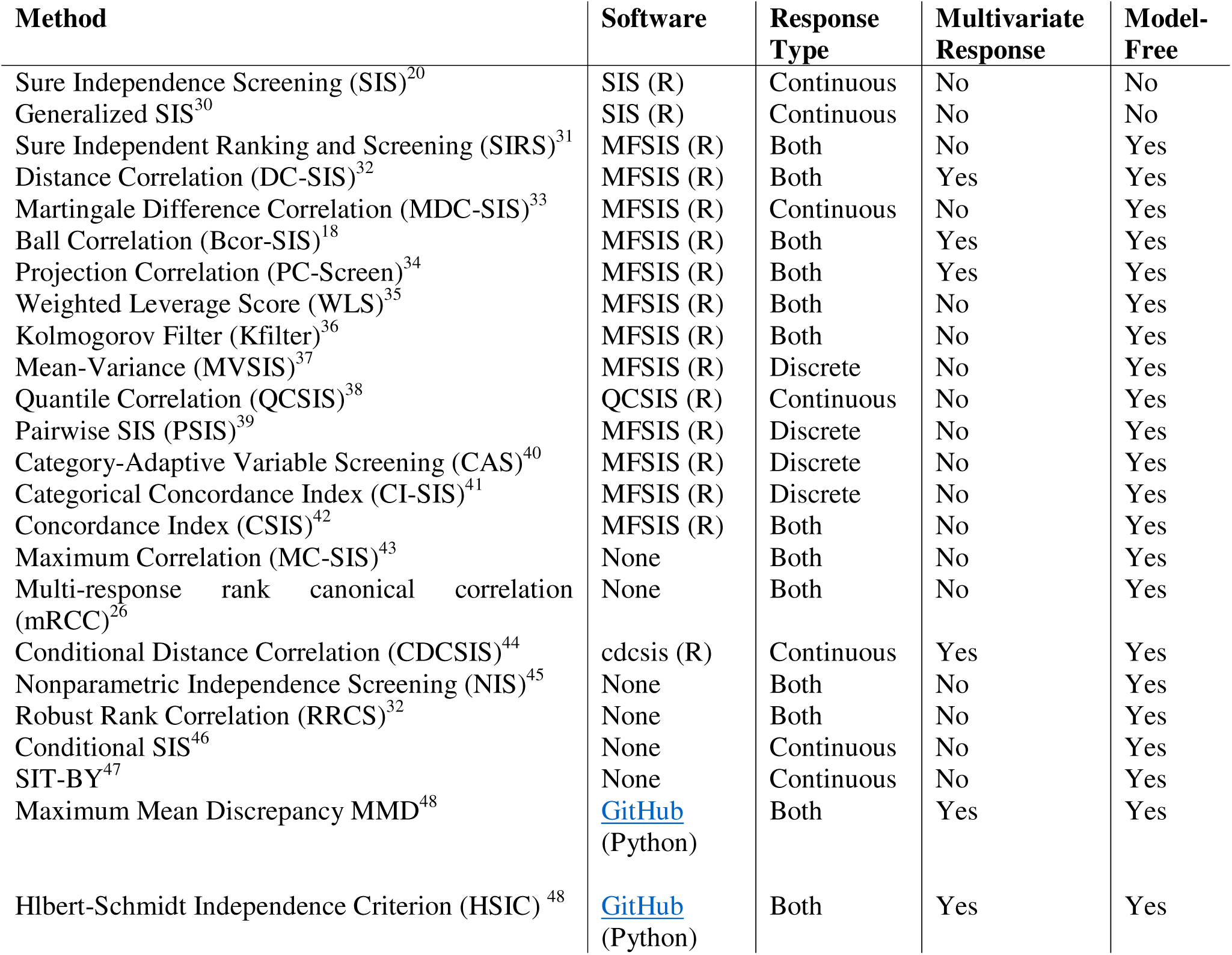
Summary of Sure Screening Methods.

A common trend in the sure screening literature is constructing iterative procedures to improve feature selection when collinearity, or more general associations, between unimportant and important features exist. One can imagine that many unimportant predictors may be marginally correlated with the response through association with another important feature. These procedures often attempt to screen out these unimportant features in a second screening stage. The common structure of these iterative procedures is best demonstrated by the original sure independent screening (SIS) paper^20^. The authors propose a three stage procedure where SIS is used to initially shrink the feature set to d_1_, followed by some stage 2 model selection method like smoothly clipped absolute deviation (SCAD)^28^ regression or the least absolute shrinkage and selection operator (LASSO)^29^. Finally, the residuals are treated as the new response, and SIS is applied to the remaining p- d_1_ features to arrive at the final model. While many authors provide the framework for a similar iterative procedure to improve their proposed screening strategy, rarely are the iterative procedures available in software, so specific publications must be referenced for implementation guidelines.

### Selecting the Model Size

An important question in the broader scope of feature selection is how to choose the model size d. Some authors and existing software suggest selecting d based on sample size. For instance, initial guidelines suggested that one should screen the feature set down to size *d*_1_ = *n*/ *log* (*n*) *or d*_1_ =*n* - 1^20^. Recent attempts at selecting the model size generally fall into two camps: direct threshold selection, and two stage approaches, with most authors suggesting a two-stage approach. Regardless of which general framework is used, false discovery rate (FDR) control is the most common approach to arrive at a final model size ^34,47–52^. While FDR control has been studied in more classical settings when p-values are available ^53–55^, general methods for FDR-motivated feature screening have only been recently proposed^56^.

Initially, a one-step filter with FDR control was proposed. This method is applicable to linear models with general design matrices^56^, though it’s limited to low dimensional problems (*p* ≤ *n*/2). To address that restriction, the model-X knockoff filter was introduced^57^ which provides FDR control for high dimensional problems when the conditional distribution of the response is both unknown and arbitrary. However, the theoretical results require that the distribution of the covariates is known. While they provide preliminary supporting evidence that the procedure can be reasonably applied when this distribution is unknown or estimated, there is still question about the quality of FDR control when the assumptions are violated^58^. Another limitation of purely knockoff-based filters is they generally do not possess the sure screening property. More recently, there has been some interest in data-splitting as a strategy for controlling FDR for general screening methods without the use of knockoffs. Xu Guo et al^50^ propose the REflection via Data Splitting method, which, under some assumptions, can perform FDR based threshold selection for any nonzero “utility statistic”, while maintaining the sure screening property. In this context, a “utility statistic” is a non-zero measure of association between a feature and the response which has the sure screening property. The authors demonstrate how this procedure can be applied to several previously proposed sure screening methods including correlation screening, rank-correlation screening, and Kolmogorov filter screening. Notably, applying this procedure to new utility statistics requires analytic derivation of a tuning parameter, which must satisfy the assumptions of the procedure.

The second class of methods, i.e. two-stage approaches, use a screening procedure to shrink the feature set to a size that satisfies the assumptions of more a powerful second stage method, which selects the final number of features in a data driven fashion. For instance, work has been done to integrate the knockoff framework with sure screening methods. Specifically, a two-stage procedure has been proposed, where the data are initially split into two sets, say, A and B. Without loss of generality, screening is performed on set A in stage 1 to select (*d*_1_ ≤ *n*/2) features, followed by applying the knockoff filter for FDR control proposed previously^56^ on the screened feature set using set B^59^. The authors discuss how the stage one screening procedure can be implemented as a sure screening procedure. Since then, some authors have started to incorporate this “screen-then-control” strategy for introducing some FDR control in the pipeline for screening methods with more relaxed assumptions about the data and model ^34,48,49^.

Unfortunately to these authors’ current knowledge, general software does not exist for FDR controlled feature screening for either the knockoff or data splitting strategies. However, readers interested in FDR control can pair existing sure screening software with the R package “knockoff” to build a 2-stage screening procedure as described above.

### Cross-Validated Screening

When developing machine learning models, one may be worried about over fitting. This can be alleviated using a cross-validation (CV) framework, where features can be screened within each fold and results are then bagged across folds, only selecting those features which were selected in a majority of the folds. Using this strategy, generalizability is introduced into the screening procedure for new data. This process can be repeated across many CV repetitions, bagging across repetitions. In the omics data context, this strategy can reduce over-fitting of a screening procedure to the training set, so that features which are more predictive of the response or outcome of interest can be identified.

## Data Applications

In this section we investigate the efficacy of various screening approaches for improving the performance of standard classification models. We start with a brief simulation study where the true important features are known to demonstrate small sample performance of the considered approaches, as well as investigate the cross validated approach discussed above. We then apply the considered approaches to a series of real data omics applications and benchmark relative performance of the methods.

### Simulated Data

First, the behavior of feature screening methods in a simulated data scenario was investigated. A total of p features, where 50 are considered “important” were simulated from a multivariate normal distribution, where important features each shared a pairwise correlation of 0.2. All other features were simulated from independent normal distributions. All features were simulated with mean 0 and unit variance. Each response was simulated as a Bernoulli random variable with a mean equal to a linear combination of important features passed through a standard logistic transformation. Coefficients for the important features were fixed at 1. Sample sizes of 10, 25, 50, and 100 were considered, along with a total feature set of size 1000 and 2000. A total of 100 datasets were simulated for each scenario. Feature screening was applied on the full set with and without repeated CV as described previously.

#### Real World Data Sets

The first dataset consisted of gas chromatography-mass spectrometry-based metabolomics data from urine samples^60^, which was originally used to identify key biomarkers for new onset type 1 diabetes (T1D). These data contained case-control pairs across 32 siblings, hereafter referred to as the CNMC data. These data had 91 metabolites, and a second dataset was constructed by expanding the feature set to include all metabolite ratios based on the unique metabolite pairings. This expanded dataset, hereafter referred to as CNMC_R, consisted of 4095 total features. This data was chosen to potentially identify any interesting behavior of screening approaches on smaller feature sets, while allowing comparisons to the larger expanded feature set.

The second applied example consisted of two datasets, one from each of two types of alternative splicing events, alternative 3’ splice site (A3SS) and retained intron (RI) splice events^61^. Data was obtained from the Human Islet Research Network (HIRN). The HIRN datasets had 6618 and 4078 features for A3SS and RI sets, respectively. These data are from a case control study, with 24 healthy controls and 24 cases of new onset type 1 diabetes. The goal of the original analysis of these data was to identify biomolecules that were highly predictive of new onset T1D. This data was chosen to evaluate smaller sample sizes with a large feature space.

The third dataset was a plasma metabolomics dataset obtained from The Environmental Determinants of Diabetes in the Young (TEDDY) consortium, as described in Li et al^62^. Briefly, the TEDDY data used here are from a matched case-control study where samples were plasma samples from 3 months prior to a case developing islet autoimmunity. The TEDDY 3-month metabolomics dataset consisted of 221 case-control pairs for a total of 441 subjects and 142 measured metabolites. This dataset was chosen as our low dimensional case study with a large sample size.

#### Screening Methods

Screening methods were chosen under the requirement that a method’s code had to 1) be open-source, and 2) be implemented entirely in R. As a result, methods BcorSIS, CAS, CSIS, DCSIS, PSIS, SIRS, and WLS were considered in this study. A sequence of model size thresholds proportional to the total feature set for a given dataset (i.e. the number of retained features is *d*= *p* * *ξ* (where (takes values from 0.05 to 1 in intervals of 0.05) were considered.

The data were screened using two approaches: ordinary and cross validated. For ordinary screening, a single pass of a given screening procedure is performed on the training set for each train/test pair, and the feature set is determined for each threshold d. In the cross validated approach, each train set was split into k folds, and the feature set was selected for each threshold as described previously. This was repeated 100 times for each training set, and the final features used for training were those that occurred in more than half the repetitions.

#### Machine Learning Models

Three machine learning methods for classification were considered in each real data example. Linear support vector machines^63^ were used as they generally benefit from a more parsimonious feature space, particularly when the feature space may contain complex nonlinear relationships. The other two methods selected perform some amount of feature selection as part of the optimization procedure (i.e., embedding methods). These methods were penalized logistic regression with elastic net penalty^64^ and random forest^15^. Both urine metabolomics sets were trained using repeated 4-fold CV with 15 repeats. All models fit using the HIRN datasets were trained using 3-fold CV with 20 repetitions. Models fit using the TEDDY dataset were trained using 12-fold CV with 20 repetitions. The CV schemes were chosen to both evenly split classes across folds and allow each model to be efficiently trained in parallel. Each HIRN and the CNMC datasets were randomly split into training and testing sets 50 times, with 50% of the data assigned to the test split, and 50% in the training set. Due to the much larger sample size, the TEDDY dataset was split into training and testing sets using 90% and 10% of the data, respectively.

#### Performance Metrics

In the simulated data example, the true and false positive rates of important feature set recovery was computed across model sizes increasing from 0.05 * p to p, where p is the number of predictors in the model. Since the TPR and FPR monotonically increase as the model size increases for a screening procedure, the results were visualized in a receiver operating characteristic (ROC)-like curve that measures feature set recovery of the screening method.

For each real data application and within each dataset, individual screening methods along with cross-validated screening were evaluated using ROC area under the curve (AUC). The performance trajectory with respect to ROC AUC is reported as a function of the number of features retained. Our various datasets have different dimensionally and signal, so we use a centering strategy to benchmark performance across all datasets. We first “center” the ROC AUC performance trajectory by subtracting the ROC AUC observed when using the full feature set. This centering is performed for each dataset, model, and screening method separately. The trajectories are expressed as a function of the proportion of features retained rather than the absolute amount, and we fit a Gaussian process with Matern 5/2 kernel^65^ to the set of trajectories within a screening method to create, in some sense, an “average trajectory”. The computation time was also recorded for each method in seconds per 1000 features screened.

As an additional comparison, the pairwise Pearson correlation between methods, computed with respect to variable importance extracted from the highest performing random forest models, was computed. Since each screening method could retain potentially different sets of features, Pearson correlation is calculated using the entire feature set after setting the importance for each excluded feature to 0. This allowed for information about both the magnitude of importance and the retained feature set to be encoded in the pairwise correlation between methods.

## Results and Discussion

The results for the simulated data analyses are shown in Figure 1. In panel A, the recovery of the important feature set is visualized as an ROC-like curve, where TPR is the proportion of recovered features that were important, and FPR is the proportion of recovered features that were not important. Panel B shows the corresponding runtimes for each method. As expected, each screening method improved as the sample size increases, reflecting the asymptotic properties of sure screening. There was little difference between cross validated and ordinary approaches. While a testing set was not present in this simulated example, CV did not lower the performance of any particular screening approach. Additionally, CSIS and DCSIS had a noticeably longer runtime than the other approaches considered. As presented below, this was also reflected in the real data applications. Notably, there was near perfect recovery of the important feature set, while minimizing false discovery, with 100 samples, and the improvement in AUC as a function of sample size is reduced for sample sizes of at least 50, as shown in Supplementary Figure 1. While not all omics studies have the luxury of 50 samples, it is still an attainable number for some larger analyses.

**Fig 1.**
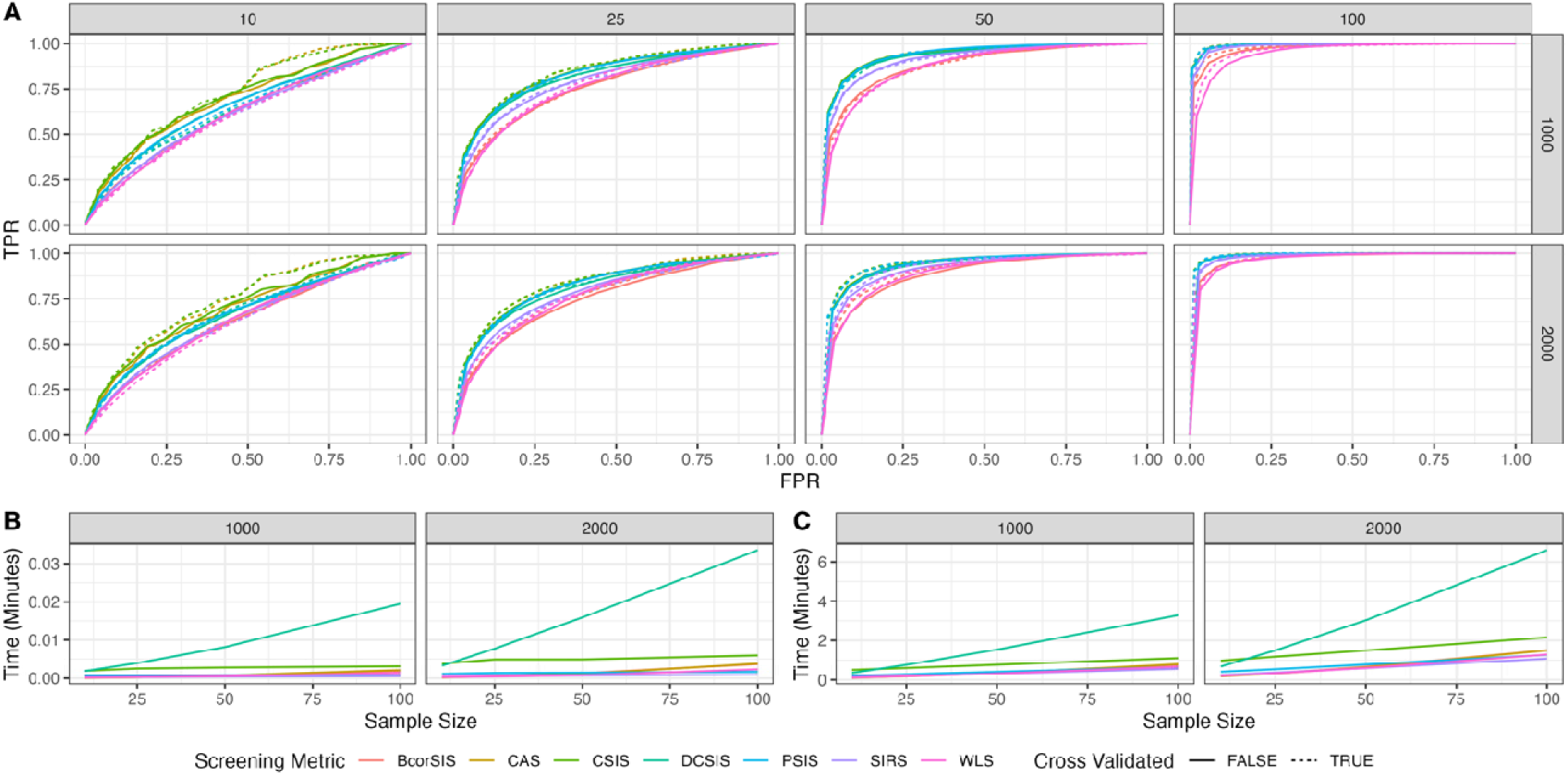
**Normal Simulation Study Results**: Results for “important” feature set recovery in th simulated data example. In Panel **A**, ROC-like curves are displayed for ordinary and cross-validated approaches, which capture a methods ability to recover the true important feature set. Panels **B** and **C** show the computation time of each method in minutes for the ordinary and cross-validated approaches, respectively.

Figures 2 and 3 give the classification performance with respect to ROC AUC as function of the number of features retained after screening for the CMNC and CMNC_R datasets corresponding to the urine metabolomics dataset. Across both datasets, the linear SVM benefited the most from feature screening, with respect to test performance. This makes sense given the underlying optimization problem linear SVM attempts to solve. Penalized logistic regression benefited from general feature screening when provided a larger initial feature set as in the CNMC_R data, compared to the original dataset with only 91 features. Random forest performed well regardless of the number of features screened out. Since random forest performs some implicit feature selection, this implies that random forest did a good job at picking out signal in these datasets without requiring the help of stage 1 feature screening. Notably, there was a visible difference in the AUC trajectory for the training set between ordinary and cross validated approaches. Specifically, the training curves were generally lower for the CV approach compared to the ordinary approach, with little difference in test performance. This indicated that CV was, to some extent, preventing the feature selection procedure from overfitting the training set. There was little difference between individual screening methods for random forest, though more separation was seen for linear SVM, and logistic regression fell in the middle.

**Fig 2.**
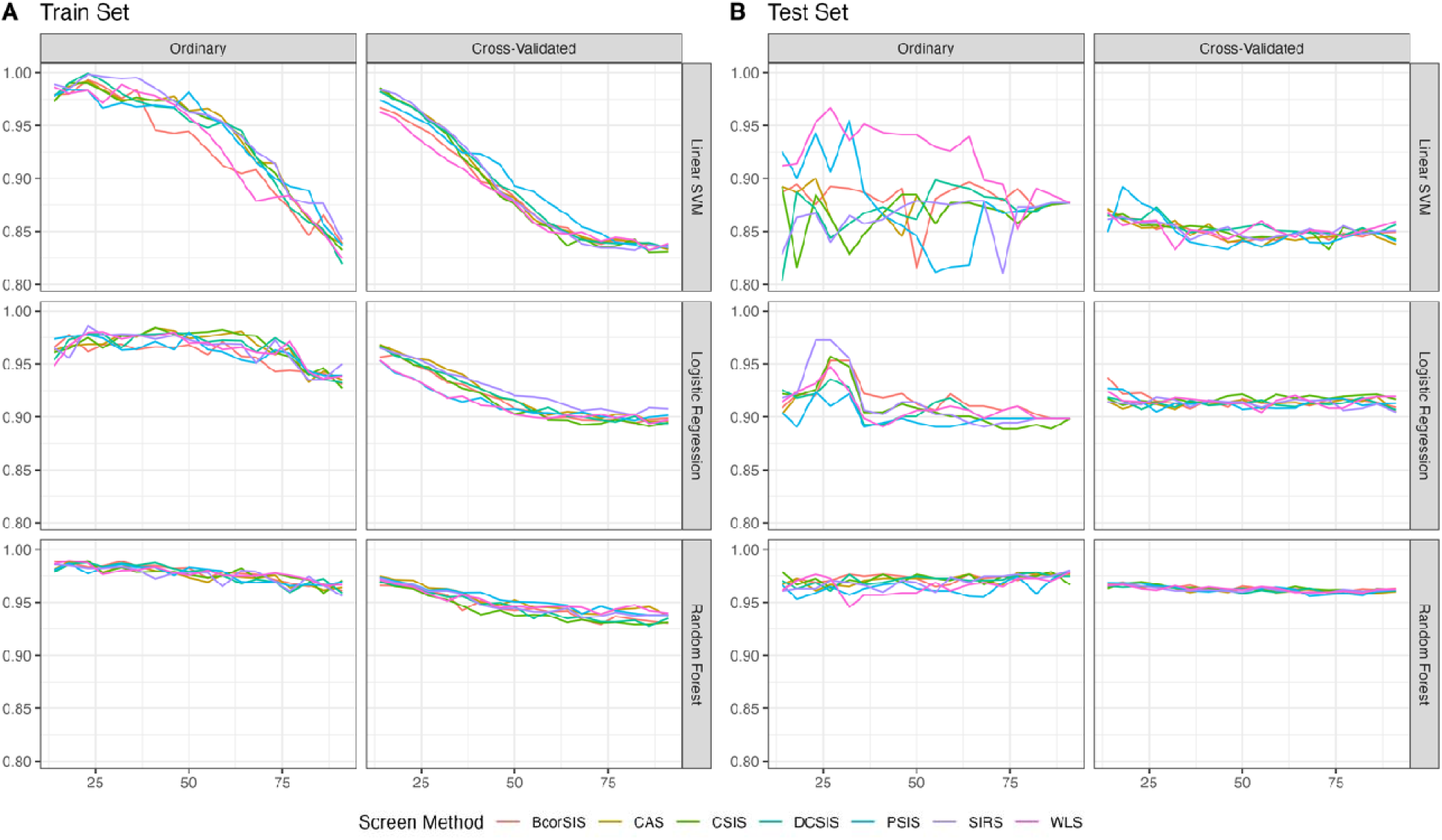
**Urine Metabolomics Application Results - CNMC**: Various feature screening methods applied to the CNMC urine metabolomics data. Panel **A** shows each classifier’s performance with respect to ROC AUC as a function of the number of features retained by the screening procedures. Similarly, panel **B** shows the test set performance.

**Fig 3.**
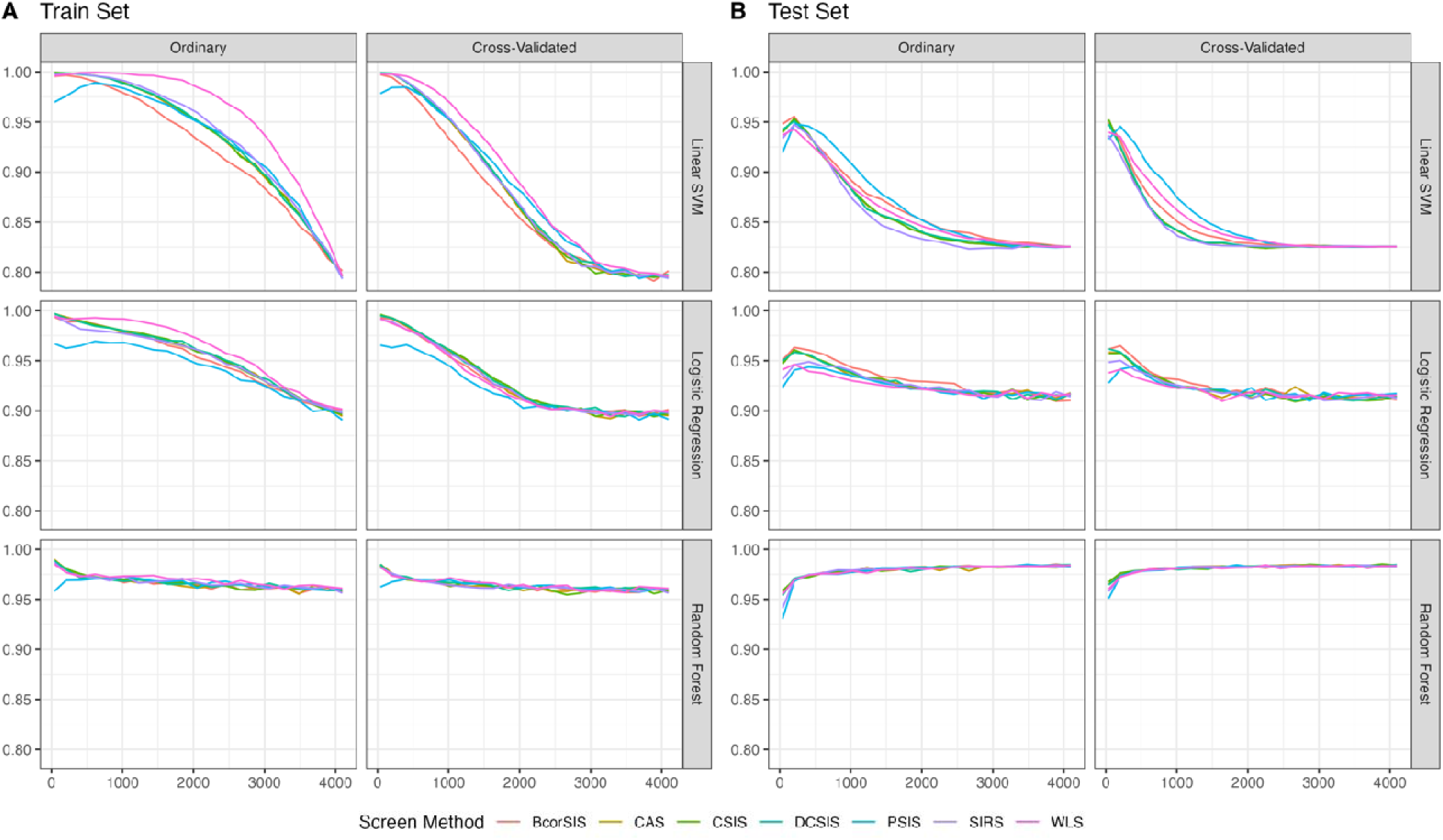
**Urine Metabolomics Application Results – CNMC_R**: Various feature screening methods applied to the CNMC_R urine metabolomics data. Panel **A** shows each classifiers performance with respect to ROC AUC as a function of the number of features retained by the screening procedures. Similarly, panel **B** shows the test set performance.

The results for the HIRN applications are displayed in Figures 4 and 5. For both datasets, poor behavior was seen for the CAS method, particularly as the number of kept features shrunk towards zero. This issue is less evident in the RI dataset, though still present, which may indicate it screened out important features too early on. Every other method had similar performance, except for WLS and PSIS showing slightly worse behavior as the feature sets got close to 0. As opposed to the CMNC datasets, there is not a clear improvement of ROC AUC for any workflow fit using either HIRN dataset. That said, the most workflows yield consistent performance with the full feature set, and which typically only decreased when the number of retained features approached zero. This is somewhat unsurprising, as we knew a-priori that the A3SS set had very weak signal, and the RI set had very strong signal, which are both situations where a screening strategy is unlikely to benefit predictive modeling.

**Fig 4.**
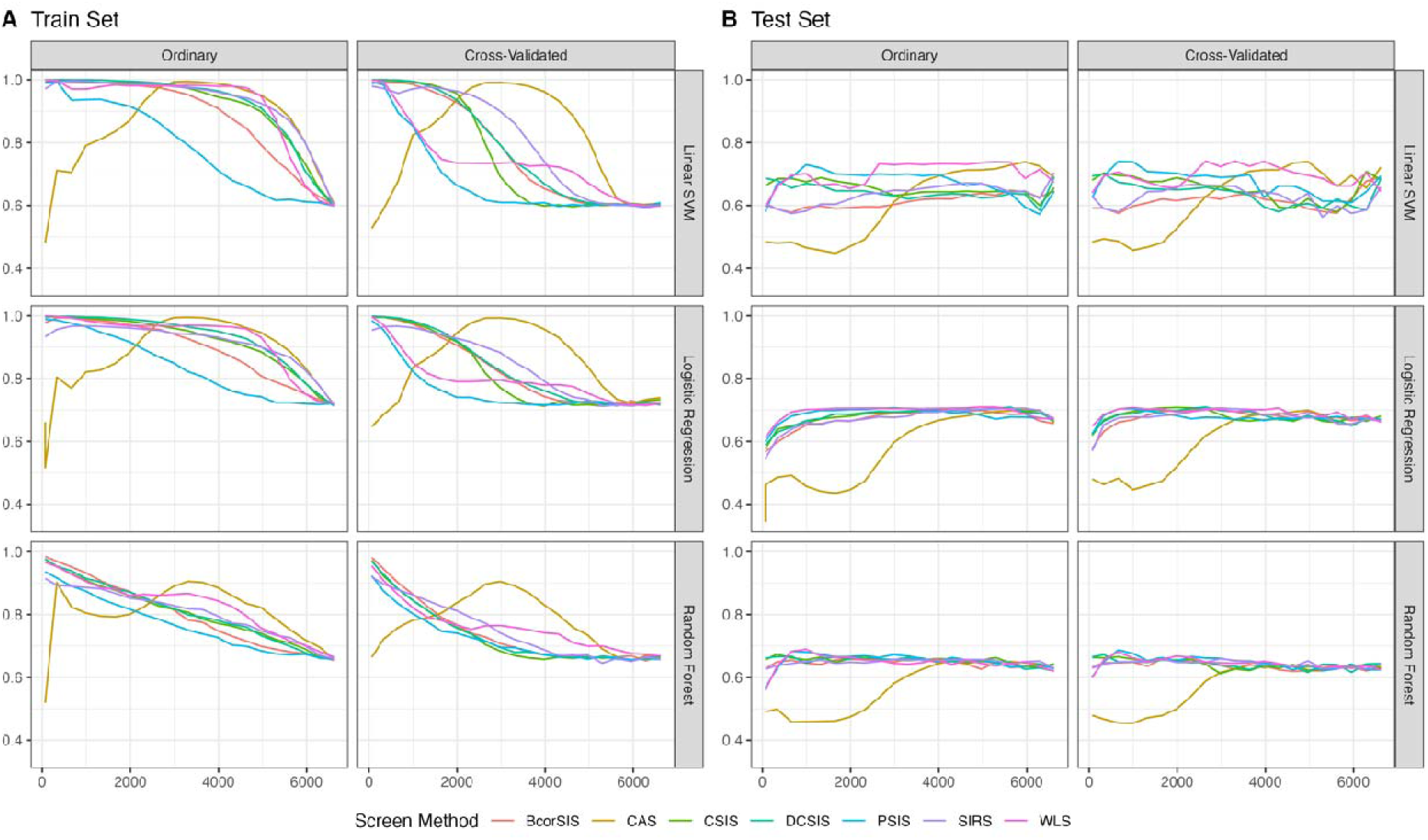
**HIRN Application Results – A3SS**: Various feature screening methods applied to th HIRN A3SS data. Panel **A** shows each classifiers performance with respect to ROC AUC as function of the number of features retained by the screening procedures. Similarly, panel **B** shows the test set performance.

**Fig 5.**
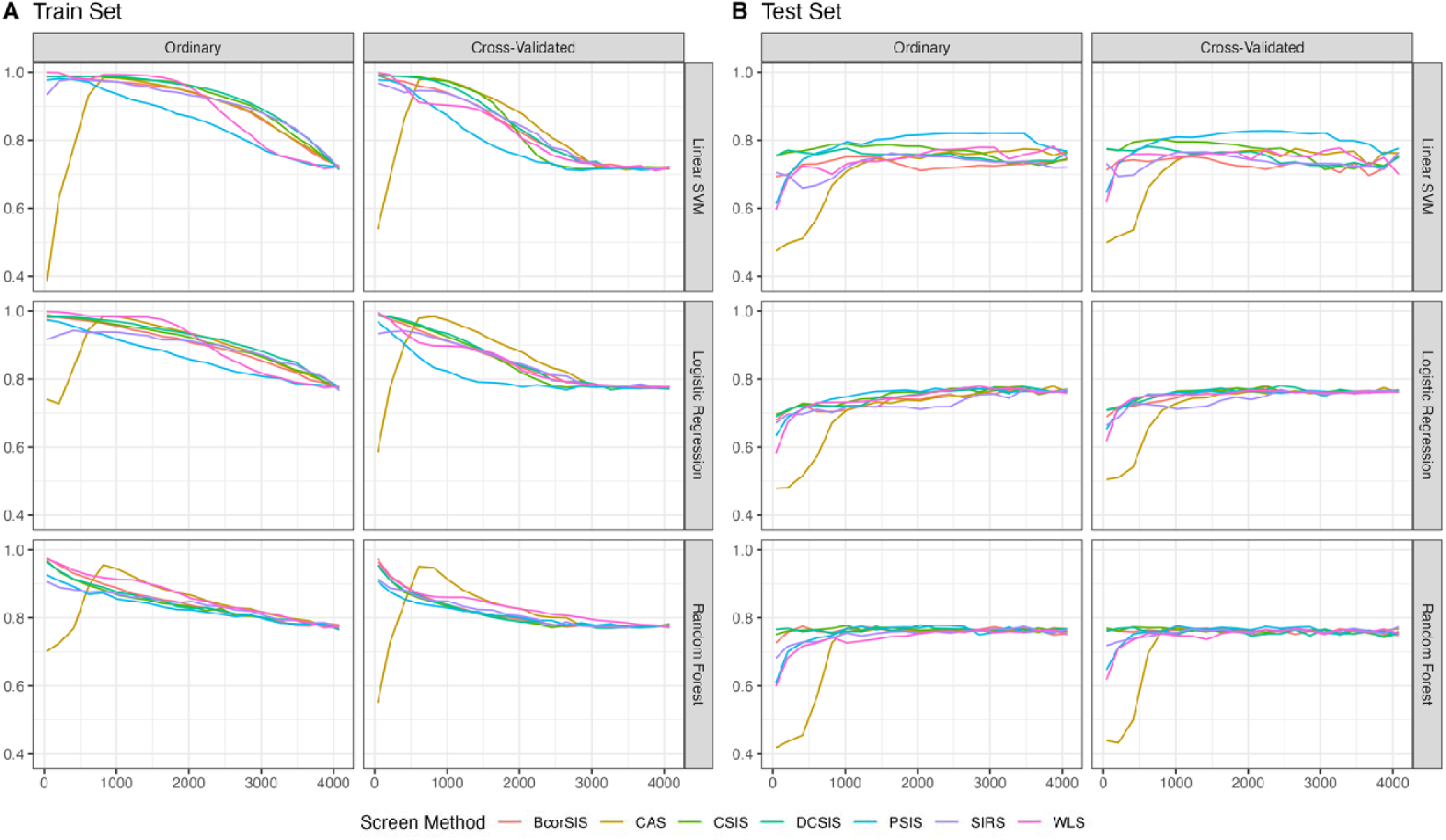
**HIRN Application Results – RI**: Various feature screening methods applied to the HIRN RI data. Panel **A** shows each classifiers performance with respect to ROC AUC as a function of the number of features retained by the screening procedures. Similarly, panel **B** shows the test set performance.

TEDDY analysis results are displayed in Figure 6. Logistic regression and SVM benefit and see a large improvement in ROC AUC as the number of retained features shrinks, and there is noticeable separation between the applied screening methods. Additionally, random forest benefits from several feature selection methods considered in both the ordinary and cross validated cases, without a large decrease in performance as the number of retained features approaches 0. As with the previous datasets, the cross-validation approach results in similar performance in the testing set, and a delayed increase to peak training set performance.

**Fig 6.**
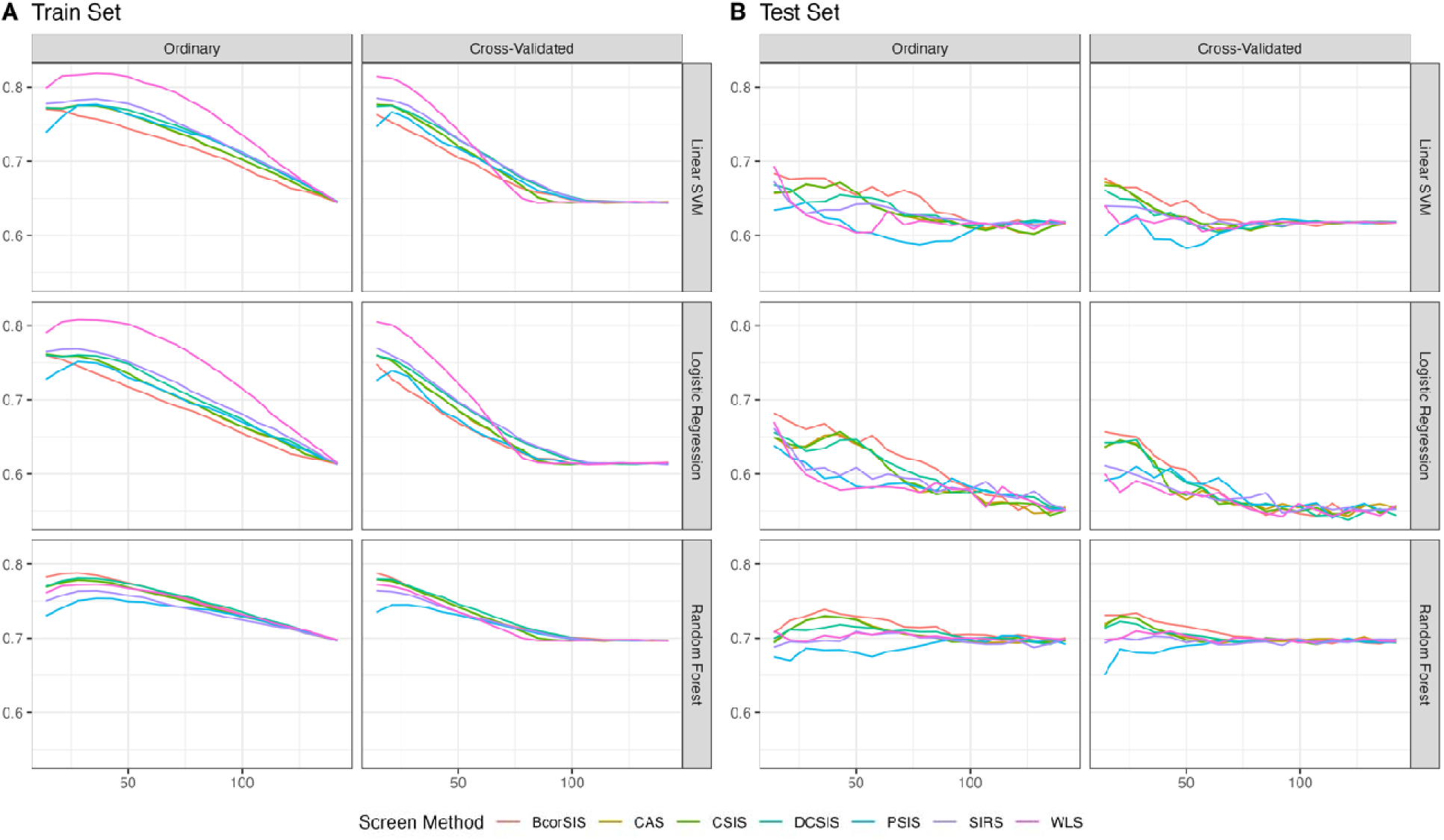
TEDDY Plasma Metabolomics Application Results –. Various feature screening methods applied to the TEDDY metabolomics data. Panel **A** shows each classifiers performance with respect to ROC AUC as a function of the number of features retained by the screening procedures. Similarly, panel **B** shows the test set performance.

The fitted Gaussian process regressions are displayed in Figure 8, along with the computation time for each screening method. For most screening methods and on average across datasets, we see improved predictive performance for models trained using the retained featur set compared to the model trained using the full feature set, until the retained feature set is roughly 10% of the full feature set. In that case, we see a large decrease in relative predictive performance if the number of retained featured gets too close to 0. That said, BcorSIS, CSIS, and DCSIS consistently outperform the others regardless of the number of retained features and also see improved performance for the smallest possible feature set size. Additionally, WLS, PSIS, and SIRS perform comparably to one another, with CAS often performing much worse than if no screening was applied. Out of the three top performing methods, BcorSIS has the shortest computation time, with CSIS and DCSIS having the slowest computation times.

In Figure 7, the Pearson correlation of mean variable importance across screening methods is shown. The mean was taken across replicates and feature sets, and feature importance was extracted from random forest models. Overall, the correlation patterns displayed agree with the trends in predictive performance discussed above. That is, the best performing methods are generally more correlated with runner ups, and not as correlated with the lowest performing methods. For instance, in the A3SS and RI datasets, the CAS method had much lower correlation with all other methods. This indicated that poor predictive performance on those data was potentially due to CAS retrieving a different, less predictive feature set. That said, there are some additional interesting observations. For the RI data, cross validated screening increased the correlation of CAS with other methods, indicating that cross validated screening can help improve the general predictive performance of the recovered feature set. Note, this same trend was not seen in the A3SS dataset, potentially due to its low signal to noise ratio *a priori*, as was seen previously^66^. Additionally, there is a similar blocked pattern in correlation matrices for both the urine metabolomics data, and the TEDDY metabolomics data, which indicates certain screening methods can potentially exhibit similar behavior across various -omics types.

**Fig 7.**
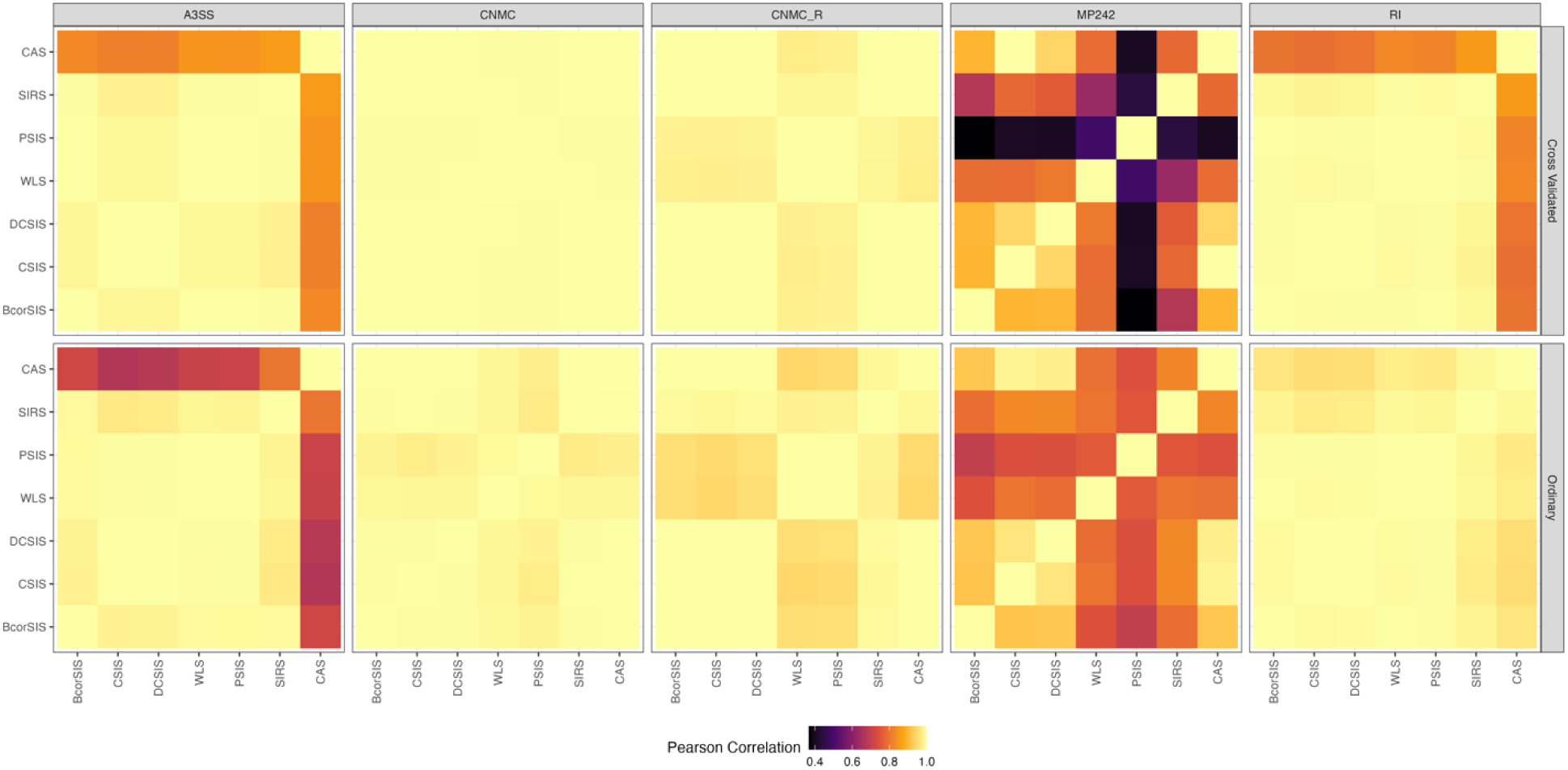
**Variable Importance Comparison**: The correlation of mean variable importance extracted from random forest models between each screening metho testing. We separate the results by dataset and if cross validated screening were applied or not.

**Fig 8.**
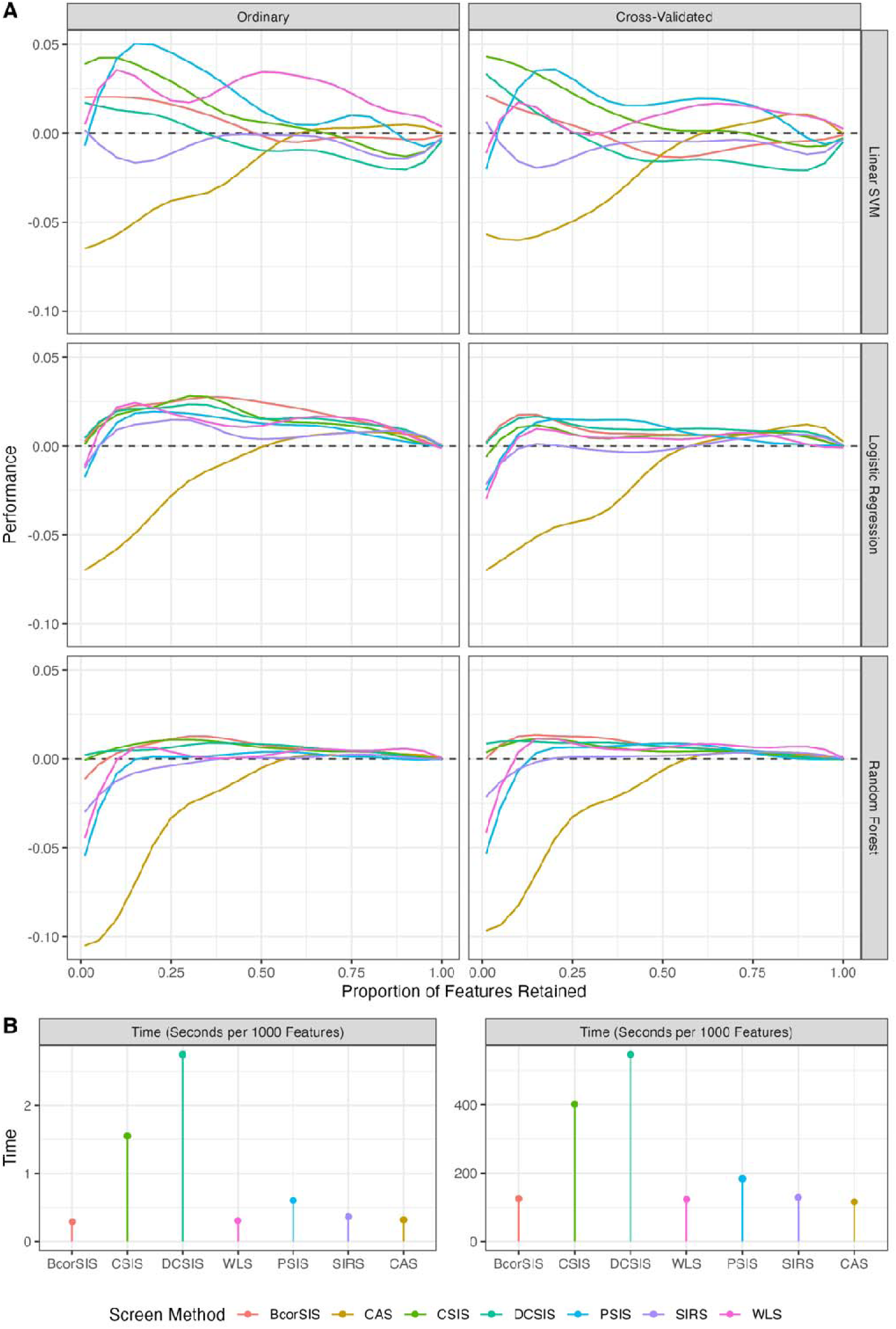
Overall Comparison: Panel **A** shows the GP regression fit across various real data applications for each screening method. Panel **B** shows the computation time in seconds per 1000 features screened for each screening method. In both panels, results for the single pass method are displayed on the left, and results for the cross validated method are displayed on the right.

## Conclusion

Often, there is much to consider when choosing a feature selection strategy. Thus, the primary goal of this paper is to inform and provide tools to practitioners who are interested in screening approaches with fewer assumptions than the filter methods common in the omics space. There are several situations where feature screening should be considered. The primary use case is when the feature set is overwhelmingly large, and it becomes infeasible to pass the full feature set into a more sophisticated feature selection method. In this case, practitioners should consider adopting the multi-stage strategies common in the feature screening literature, where an initial pass of feature screening is performed before passing the reduced feature set into a more sophisticated method. However, if the analysis timeline needs to be expedited, feature screening can be used as a one-step feature selection procedure that, at worse, reduces the feature set and run-time of the modeling task while maintaining performance compared to the full feature set. Though some wrappers (e.g. ROFI) and embedders (e.g. LASSO) do have methods for reducing the amount of over-fitting to the training set, a simple cross-validation strategy that can also reduce overfitting for any screening metrics was presented and investigated here. This can be performed as an initial screening strategy to significantly reduce the computation time of a second stage strategy like ROFI.

While the results here show varying performance depending on the data application, when it comes to sure independence screening in omics data, we identified three methods that consistently overperform across datasets: BcorSIS, DCSIS, and CSIS. However, CSIS and DCSIS have much longer run-times than BcorSIS, which suggests BcorSIS may be a good fit for general use. Additionally, it was shown that CAS consistently underperforms in these applications, often yielding a model that has lower predictive performance than one trained on the full feature set. In general, analyses should be tailored to specific contexts. We suggest that practitioners contextualize any screening method, or feature selection method at large, with the analysis problem at hand. To assist with this, the key properties and available software for each model free screening method is available in Table 1, along with the relevant citations for those interested in further investigation.

As with any review and broad application of methods, there are limitations to this study. First and foremost, data driven insights into the multi-stage iterative procedures individual screening methods typically propose to improve were not provided. This is closely tied to the idea of FDR control, which is a relatively new concept in the field, which was also omitted in the real data applications. However, as the field progresses, it is expected that more software will become available for FDR motivated screening approaches, which provides an interesting avenue of future research. Additionally, missing data is a common problem in certain omics applications, specifically proteomics. While the real data applications had minimal missing data, there has been some work on the impacts of missing data and imputation on sure screening procedures ^67^. Further research is required for those specifically interested in shotgun proteomics. Lastly, theoretical results, which we confirm in simulation, state that the sure screening property is asymptotic with sample size. Modern -omics analysis typically have limited samples, but as technology (i.e. multiplexing) improves sample sizes in high throughput omics, feature screening will become an increasingly viable option. Overall, this manuscript aimed to provide practitioners with knowledge of more sophisticated methods for feature screening, so that they can be more equipped to handle the feature selection problem in omics data analysis.

## Data Availability

The CMNC dataset is available at https://data.pnnl.gov/group/nodes/dataset/34049. The HIRN dataset is available at https://data.pnnl.gov/group/nodes/dataset/33494. The TEDDY dataset can be requested via the TEDDY access page (https://teddy.epi.usf.edu/research/). The availability of data is subject to approval by the TEDDY Ancillary Studies Committee.

## Acknowledgements

This work was support in part by National Institutes of Health contract HHSN267200700014C and grants R01 DK138355 (ESN, BJMWR, TOM) and U01 KD127786-S1 (BJMWR) from the National Institute of Diabetes and Digestive and Kidney Disease. Mass spectrometry analyses were performed in the Environmental and Molecular Sciences Laboratory, a national scientific user facility sponsored by the U.S. Department of Energy (DOE) and located on the campus of PNNL in Richland, WA. PNNL is operated by Battelle Memorial Institute for the DOE under contract DEAC05-76RLO1830.

The TEDDY Study is funded by U01 DK63829, U01 DK63861, U01 DK63821, U01 DK63865, U01 DK63863, U01 DK63836, U01 DK63790, UC4 DK63829, UC4 DK63861, UC4 DK63821, UC4 DK63865, UC4 DK63863, UC4 DK63836, UC4 DK95300, UC4 DK100238, UC4 DK106955, UC4 DK112243, UC4 DK117483, U01 DK124166, U01 DK128847, and Contract No. HHSN267200700014C from the National Institute of Diabetes and Digestive and Kidney Diseases (NIDDK), National Institute of Allergy and Infectious Diseases (NIAID), Eunice Kennedy Shriver National Institute of Child Health and Human Development (NICHD), National Institute of Environmental Health Sciences (NIEHS), Centers for Disease Control and Prevention (CDC), and Breakthrough T1D (formerly JDRF). This work is supported in part by the NIH/NCATS Clinical and Translational Science Awards to the University of Florida (UL1 TR000064) and the University of Colorado (UL1 TR002535). The content is solely the responsibility of the authors and does not necessarily represent the official views of the National Institutes of Health.

## Author Contributions

EV: methodology, software, formal analysis, writing – original draft LB: conceptualization, methodology, writing - original draft JF: writing – review and editing EN: writing – review and editing, funding acquisition TM: writing – review and editing, funding acquisition BJWR: conceptualization, writing – review and editing, supervision, funding acquisition

## Notes

### Competing Interest Statement

The authors have declared no competing interest.

