## Supplemental Material for "A Benchmarking Study of Feature Screening Approaches Across Omics Classification Settings"

### Supplementary Information

**Supplementary Fig 1. Simulated Example AUC:**  The AUC for feature recovery in the simulated data example is displayed as a function of sample size.

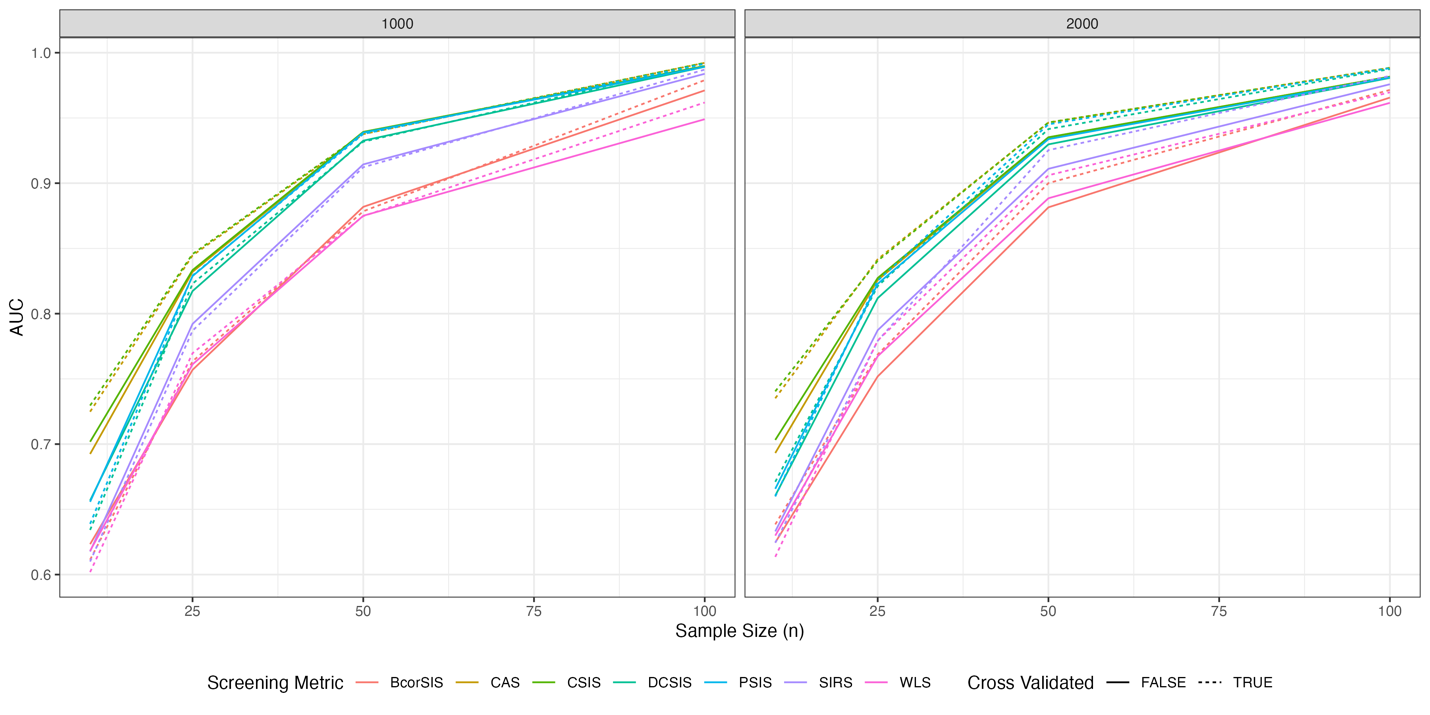
